# The spatiotemporal dynamics of semantic integration in the human brain

**DOI:** 10.1101/2022.09.02.506386

**Authors:** Elliot Murphy, Kiefer J. Forseth, Cristian Donos, Patrick S. Rollo, Nitin Tandon

## Abstract

Language depends critically on the integration of lexical information to derive semantic concepts. The neurobiology of this fundamental aspect of human cognition remains mostly unknown. Limitations of spatiotemporal resolution have previously rendered it difficult to disentangle processes involved in semantic integration from independent computations such as tracking word meaning and establishing referents. We utilized intracranial recordings (n = 58) during the presentation of orthographic descriptions that were either referential or non-referential to a common object. Referential contexts enabled high frequency broadband gamma activation (70–150 Hz) of a distributed network including the inferior frontal sulcus (IFS), medial parietal cortex (MPC) and medial temporal lobe (MTL) in the left, language-dominant hemisphere. Components of this network (IFS, MPC), alongside posterior superior temporal sulcus, showed greater engagement for trials that did not progressively isolate a limited set of referents, relative to trials that did. IFS and posterior middle temporal gyrus activity was modulated by semantic coherence, regardless of whether the sentence picked out a common object. Centered around IFS and spreading out dorsally towards middle frontal gyrus and ventrally towards anterior/posterior inferior frontal gyrus, we uncovered a cortical mosaic of functional specialization for reference, coherence and semantic narrowing. Early processing windows implicated IFS in all aspects of semantics, and in later windows sub-portions of IFS exposed a clearer functional tessellation with some remaining overlap. This work unveils how lateral fronto-temporal regions make distinctly rich semantic contributions and are involved jointly in semantic integration and conflict resolution, depending on the sub-region and processing stage.

## Introduction

Retrieving specific words from memory to refer to objects in the world is an essential contribution made by language to our cognitive abilities (Abbott, 2008; Burge, 1973; Chomsky, 2000; Davidson, 1977; Hinzen, 2007; Jackendoff, 2002; Kripke, 1980, 2013; Murphy, 2016, 2020; Pessin, 2016; Pietroski, 2018; Russell, 1905; Strawson, 1950). The fundamentally generative nature of language enables us to derive concepts using cues we may never have encountered, such as when the phrase “an elongated yellow fruit” is immediately recognized as referring to a banana. Such semantic derivations depend upon the rapid integration of multiple lexical objects into a larger structure, but this crucially occurs alongside a range of independent cognitive processes pertaining to the tracking of individual words (Berwick and Stabler, 2019; Murphy and Leivada, 2022). We know broadly that these processing engage the posterior temporal lobe (Brennan et al., 2016; Matchin and Hickok, 2020) and prefrontal and parietal cortices (Brodbeck and Pylkkänen, 2017; Ferstl et al., 2008; McMillan et al., 2012; Nieuwland et al., 2007), but there is no general consensus about the neurobiology of distinct semantic processes. Given the very rapid and distributed nature of these processes (Huettig et al., 2011; Tanenhaus et al., 1995), previous research into lexical access has been unable to disentangle retrieval from related computations, such as establishing individual word meaning, inferring semantic coherence, and narrowing the lexical search space over time. Much work into linguistic coherence has utilized scalp event-related potentials or fMRI (Boudewyn et al., 2012; Camblin et al., 2007; Ferstl and von Cramon, 2002; Jouen et al., 2021), which lack the fine spatiotemporal resolution needed to comprehensively map cortical responses. Under some analyses, the cortical substrates for language and semantic processing overlap (Ivanova et al., 2021; Sueoka et al., 2022), while others point to dissociability (Colvin et al., 2019).

In order to isolate sites involved in distinct semantic processes we used an orthographic sentence comprehension and linguistic reference paradigm in a large cohort of subjects undergoing intracranial electrocorticography (iEEG) (58 patients; 11,328 electrodes) (Arya, 2019; Flinker et al., 2011). We used broadband high gamma activity (BGA; 70–150 Hz), which strongly correlates with the functional MRI (fMRI) blood oxygen level-dependent (BOLD) signal, to index local cortical processing (Buzsáki, 2019; Conner et al., 2011). We comprehensively mapped BGA responses across the language-dominant cortex by presenting both congruent (Penolazzi et al., 2009) and incongruent (Obleser and Kotz, 2011) sentences.

We utilized orthographic stimuli that afford temporally precise lexical inputs and minimize integrative processes that are intrinsic to auditory or visual inputs. Patients generated common object names in response to written descriptions of variable lengths via rapid serial visual presentation (500 ms per word). The final word dictated whether or not the sentence referred to a common lexical item. We manipulated both the referential nature and the semantic coherence of our sentences, whilst varying syntactic structure and content to reduce predictability. This design allowed us to isolate the cortical dynamics of semantic convergence to a basic lexical item, alongside semantic narrowing and coherence. For example, “A person at the circus who makes you laugh” permits successful lexical access (e.g., clown). However, “A person at the circus who makes you commute” represents a *weak violation* of lexico-semantic rules, as it is coherent but does not refer to a common object. “A glass made of Wednesday” is semantically incoherent and additionally does not permit lexical access (i.e., a *strong violation*). Lastly, we collected additional norming data (n = 80) to quantify the point of ‘narrowing’ to analyze the extent of lexicon search effort, e.g., “It’s white and falls from the sky in winter” is likely to be inferred as “snow” before the final word, while other trials do not enable inference until the final word, e.g., “An object used for weighing”.

The high spatiotemporal resolution of intracranial recordings allowed us to probe: (1) word-by-word semantic integration; (2) narrowing of the search space as semantically salient information is integrated before the final word; and (3) integration of semantically coherent information regardless of the success of lexical access (i.e., non-referential sentences that exhibited weak vs. strong lexico-semantic violations). Together, these analyses enabled us to comprehensively elaborate the spatiotemporal dynamics of *semantic integration* in the service of *lexical access*.

## Results

### Behavioral performance

Individual reaction time (Fig. 1) averaged 1765 ms (SD: 738 ms) after the offset of the final word in the sentence. As expected, referential trials had significantly faster articulation reaction times than non-referential trials (paired t-test; *t*(1,83): 4.62, *p*-value: <.001).

**Figure 1:**
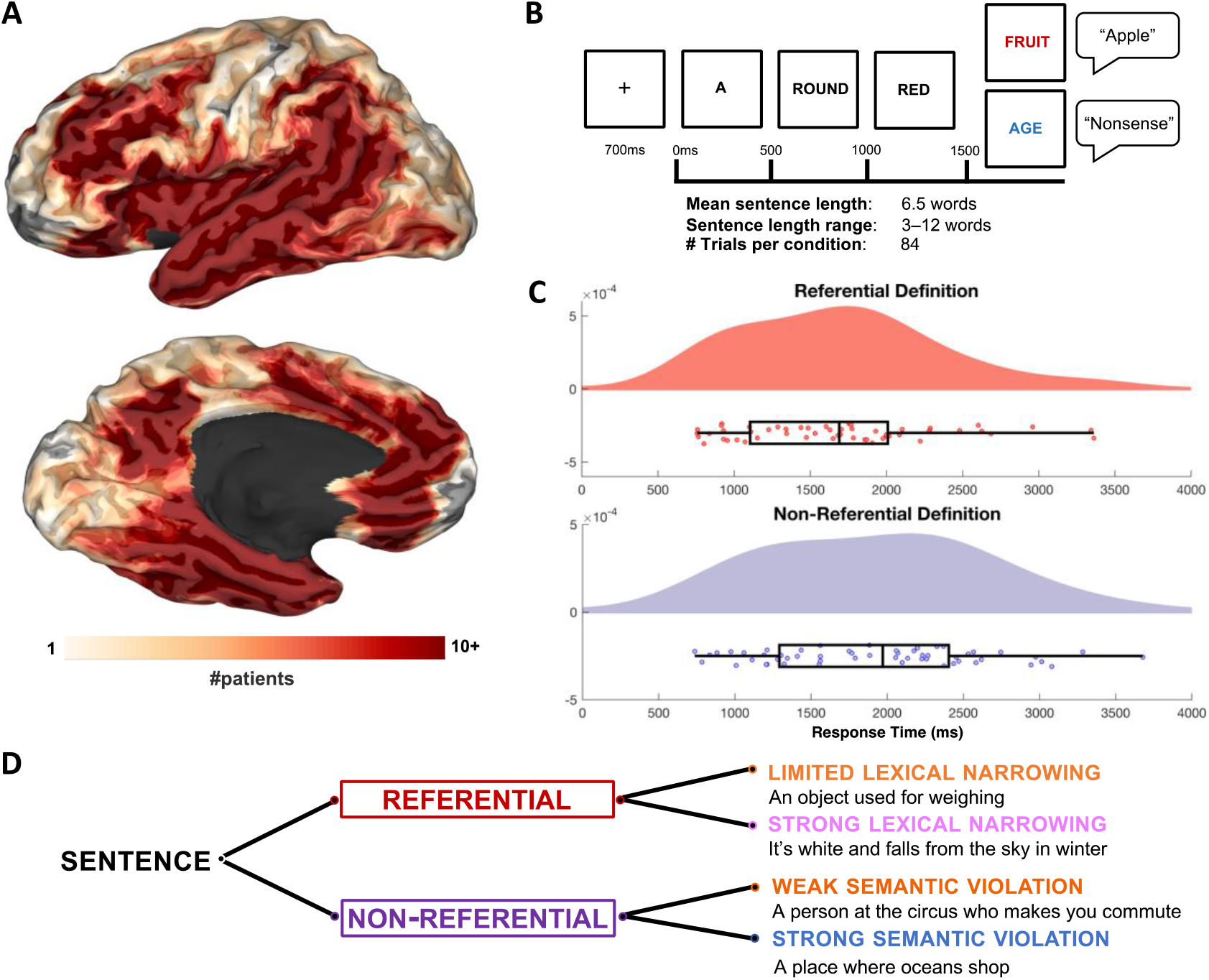
Intracranial electrode coverage with experimental paradigm. (A): Group coverage map, plotted on a semi-inflated standardized N27 surface. (B): Experimental design. Sentences were presented orthographically in rapid serial visual presentation, 500 ms per word. Patients verbally articulated their responses. (C): Time from the offset of the final word to onset of patient verbal articulation. (D): Sentence conditions split by referential status, with example trials. The three main semantic factors analyzed below that we derived from these conditions are Reference (referential vs. non-referential), Coherence (weak vs. strong semantic violation) and Narrowing (limited vs. strong lexical narrowing).

### Spatiotemporal dynamics of orthographic sentence processing

To probe the build-up of local cortical activity across successive words, we generated a population-level map of cortical activity using a surface-based mixed-effects multi-level analysis (SB-MEMA) (Conner et al., 2011; Fischl et al., 1999; Kadipasaoglu et al., 2014, 2015). This revealed serially increasing activation across the sentence duration in a distributed orthographic sentence processing network: early activation in inferior frontal gyrus (IFG), medial parietal cortex (MPC), anterior temporal lobe (ATL) and posterior middle temporal gyrus (pMTG), followed by late activation in ventro-medial prefrontal cortex (vmPFC) and posterior cingulate (Fig. 2). Three regions – the ventral temporal cortex, the inferior lateral temporo-occipital cortex and the inferior frontal sulcus (IFS) in its entire antero-posterior extent – were active throughout sentence reading and showed a clear increase in BGA at the final word and a broader spread of activity across the sentence.

**Figure 2:**
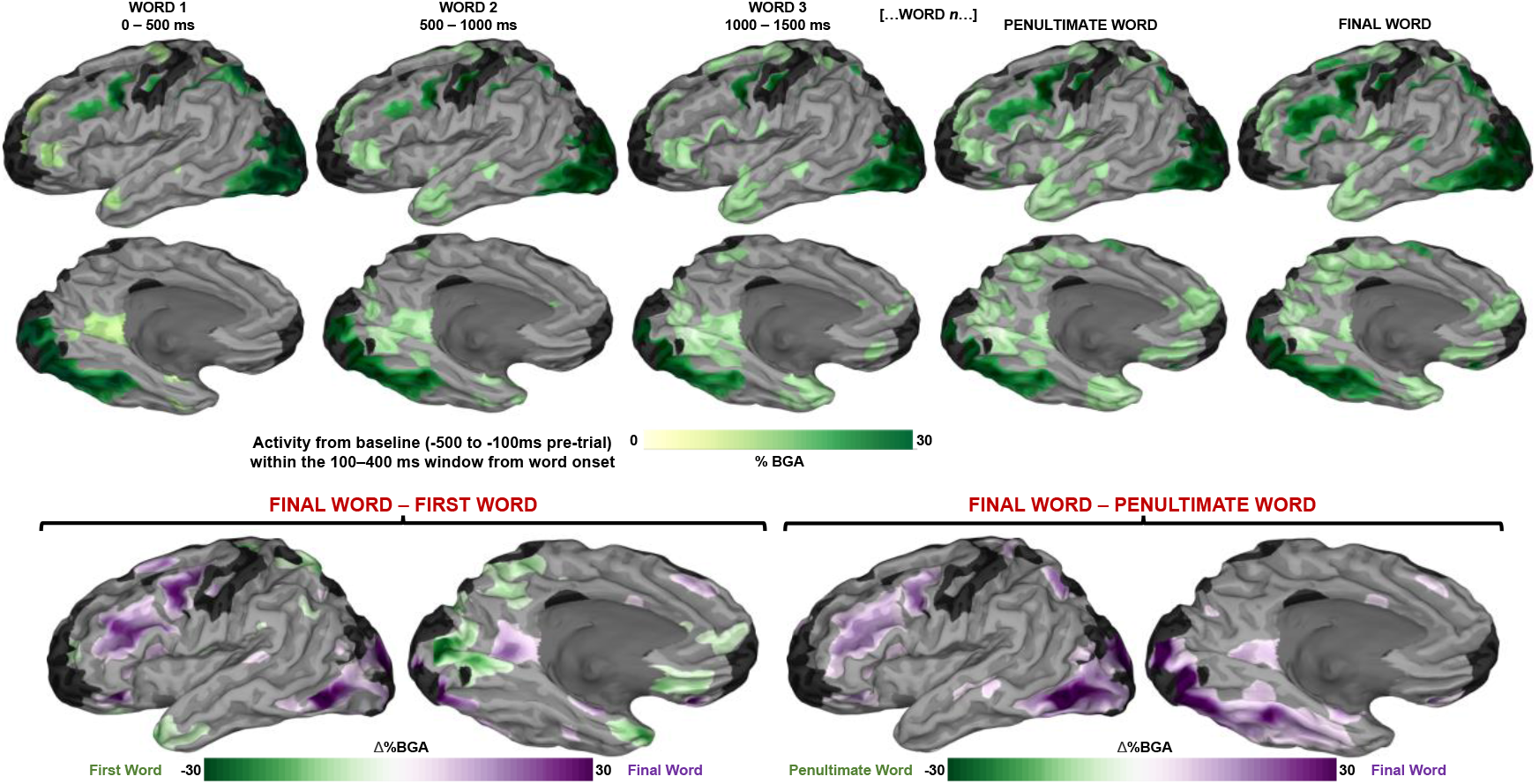
Grouped analysis for semantically coherent sentence processing. Top: Responses to orthographic stimuli (averaged across the 100–400 ms window after presentation of each word), relative to baseline (−500 ms to -100 ms before first word onset) represented using a surface-based mixed effects multi-level analysis (SB-MEMA) (thresholded at t > 1.96, patient coverage ≥ 3, corrected *p* < 0.01). Trials with only three words (2 trials) were excluded from Word 3 maps. Bottom: Contrast for final vs. first word (*left*) and final vs. penultimate word (*right*) across the same window and baseline.

### Reference to common objects

A comparison of referential vs. non-referential trials at the final word revealed greater BGA for referential trials in MFG and mid-IFS (500–700 ms: *β* = 0.08 (SD: 0.03); *p* < 0.001; 700–900 ms: *β* = 0.10 (SD: 0.05); *p* < 0.001), MPC and parahippocampal cortex (500–700 ms: *β* = 0.11 (SD: 0.06); *p* < 0.001; 700–900 ms: *β* = 0.13 (SD: 0.08); *p* < 0.001), and vmPFC (500–700 ms: *β* = 0.07 (SD: 0.01); *p* < 0.001; 700–900 ms: *β* = 0.08 (SD: 0.04); *p* < 0.001) (Fig. 3, Suppl. Fig. 1). Non-referential trials (semantically incongruous) exhibited BGA increases relative to congruous trials in posterior superior temporal cortex (500–700 ms: *β* = 0.07 (SD: 0.01); *p* < 0.001) and anterior inferior frontal gyrus (aIFG) (500–700 ms: *β* = 0.08 (SD: 0.03); *p* < 0.001) after final word onset (Fig. 3). A whole brain analysis contrasting strong and weak violations for non-referential trials showed increased activity in medial frontal cortex (300–500 ms: *β* = 0.10 (SD: 0.04); *p* = 0.001) and superior medial parietal cortex (300–500 ms: *β* = 0.09 (SD: 0.03); *p* < 0.001) for strong violations. The supero-medial parietal activations were seen in loci distinct and non-overlapping with the sites sensitive to referential vs. non-referential contrasts. Weak violations resulted in greater BGA in IFS (300–500 ms: *β* = 0.08 (SD: 0.03); *p* = 0.001), aIFG (700–900 ms: *β* = 0.11 (SD: 0.05); *p* = 0.001), angular gyrus (700–900 ms: *β* = 0.06 (SD: 0.01); *p* < 0.001) and pMTG (700–900 ms: *β* = 0.08 (SD: 0.03); *p* = 0.002).

**Figure 3:**
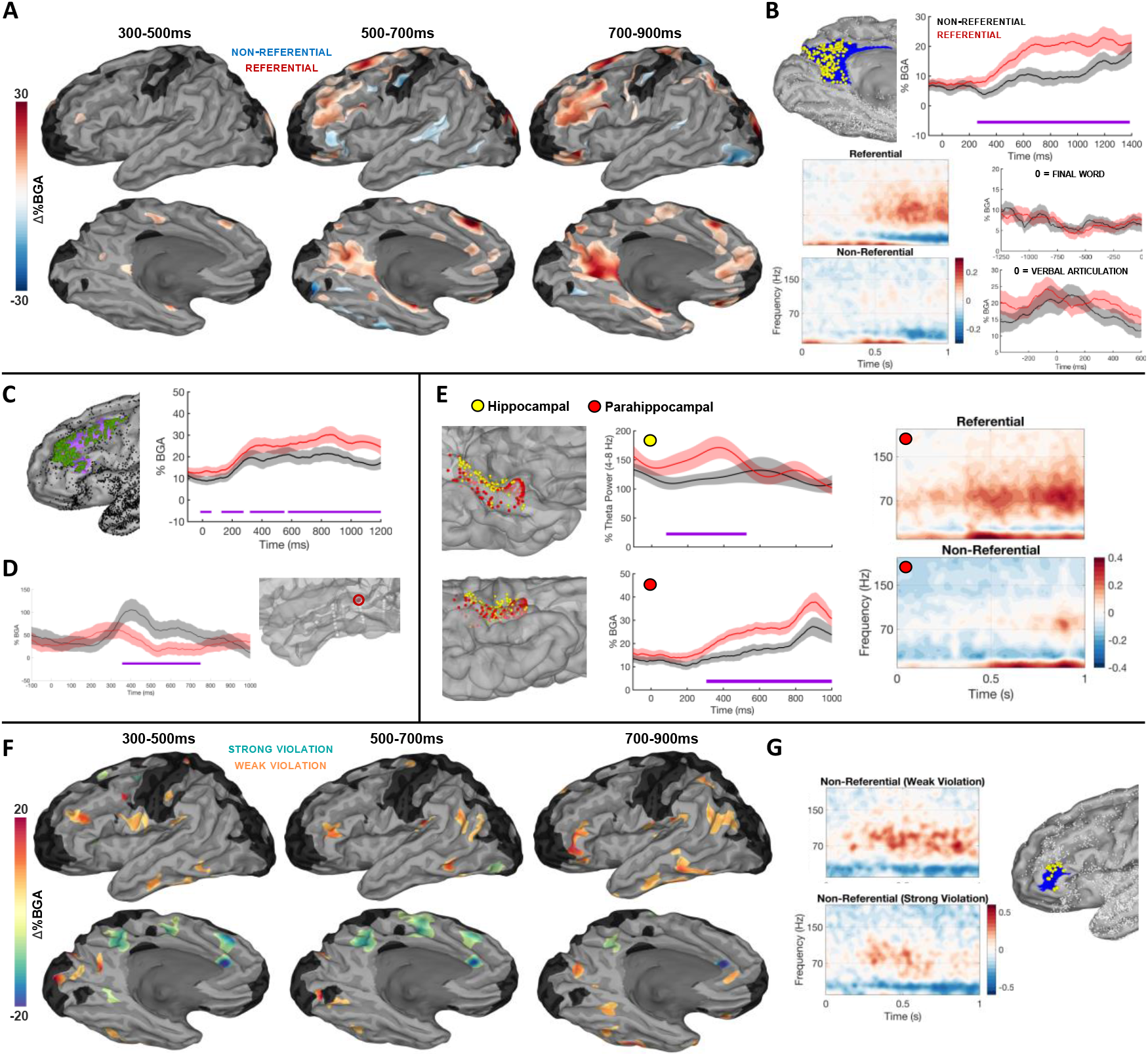
Grouped analysis for linguistic reference. (A): SB-MEMAs for referential vs non-referential trials, with red indexing greater BGA (70–150Hz) for referential sentences and blue for non-referential sentences (threshold: %BGA > 5%, t > 1.96, patient coverage ≥ 3, *p* < 0.01 corrected). Time 0 ms = onset of final word. (B): Top Left: Electrodes (yellow dots; n = 245) placed within medial parietal cortex (MPC); other electrodes are small dots. Top Right: Grouped electrode traces for BGA (red: referential; black: non-referential), error bars (colored shading) set at one standard error; significance (FDR-corrected) bars in purple. Bottom Left: Spectrographic representation of grouped electrodes in MPC time-locked to final word onset. Bottom Right: Grouped traces time-locked to pre-final word period (top) and patient verbal articulation (bottom) for MPC. (C): Grouped traces for middle frontal cortex (electrodes: 194). (D): Semantic violation effect to non-referential trials; exemplar patient with recordings from posterior superior temporal regions.(E): Left: Recording sites (334 electrodes in 42 patients) placed within either hippocampus (yellow) or parahippocampal gyrus (red) represented on a standardized pial N27 surface. Middle: Grouped representation of hippocampal theta power (top) and parahippocampal BGA (bottom) time-locked to final word onset. Right: Spectrographic representation of grouped electrodes for all active electrodes in parahippocampal cortex. (F): SB MEMAs for non-referential *weak violation* (in lexico-semantic selectional requirements and/or world knowledge) vs.non-referential *strong violation* trials, with dark orange coloration indexing greater BGA (70-150Hz) for *weak violation* sentences and turqoise coloration for *strong violation* sentences (same thresholds as (A)). (G): Electrodes and spectrograms for anterior IFG (HCP index: p47r; electrodes: 24; patients: 10) for weak vs. strong non referential trials.

### Semantic narrowing

To calibrate our stimuli, we conducted a norming study with native English speakers (n = 80) enabling the derivation of the point of *semantic narrowing*, the probability that the defined object could be identified before the presentation of the final word. Certain sentences yielded probable answers before the final word (e.g., “Something you use to *unlock* a door”), whereas other sentences were ambiguous until the final word (e.g., “What you use to measure *temperature*”). We then contrasted trials that exhibited limited semantic narrowing at the point of final word onset with those exhibiting strong semantic narrowing (Fig. 4). This was the most conservative means of isolating a contrast between limited and strong narrowing due to variability in the position of narrowing words throughout the sentence, as well as our inability to ensure that patients were specifically narrowing their definitional search at the same points as the average sentence positions in the norming data. Articulation reaction times did not differ between semantic narrowing conditions (strong narrowing: 1886 ± 708 ms; limited narrowing: 1904 ± 699 ms; paired t-test, *t*(1,34) = -0.09, *p* = .46). There was no significant difference between conditions with respect to sentence length (strong narrowing length; M: 6.8; range: 5 (4-9); limited narrowing length; M: 6.4; range: 5 (3-8); paired t-test, *t*(1,41) = 1.36, *p* = .08). There was a significant difference in the frequency of the final word between the two conditions, such that the final word in limited narrowing trials was less frequent; however, this was a small effect and is unlikely to be the primary driver of any reported BGA differences (strong narrowing final word frequency, SUBTLEXus log-frequency: 3.54; range: 4.4 (1.1-5.4); limited narrowing final word SUBTLEXus log-frequency: 3.18; range: 4.5 (0.5-5.0); paired t-test, *t*-(1,41) = 1.72; *p* = .044), and the cortical loci we document below are not typically implicated in lexical frequency sensitivity (Woolnough et al., 2021).

**Figure 4:**
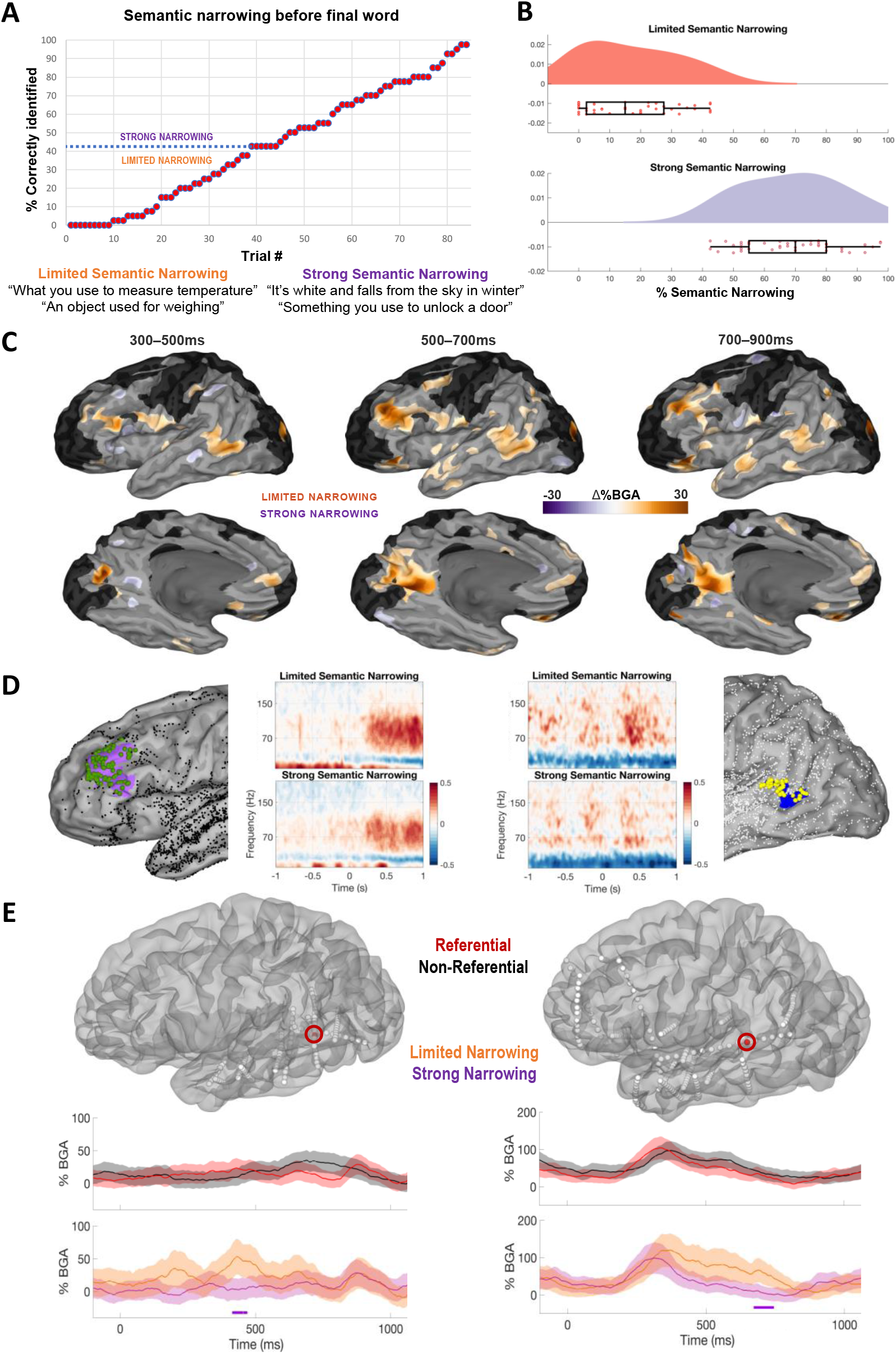
Grouped analysis for semantic narrowing in referential trials. (A): Distribution of all referential trials organized by the percentage of responses in the norming study that correctly identified the common object before the final word, indicating the cut-off point equally dividing trials into distinct conditions. Example trials are given in boxes. (B): Percentage of participants in norming study who successfully defined the common object prior to the final word, grouping all trials (1-84 referential trials) into the top and bottom half percentages to derive the two groups. Generated using Raincloud (Allen et al., 2021). (C): SB-MEMA contrasting limited narrowing (orange) and strong narrowing (purple) trials in BGA from 100-900 ms after final word onset (threshold: %BGA > 5%, t > 1.96, patient coverage ≥ 3, *p* < 0.01 corrected). (D): Left: Spectrogram and electrode placements for middle frontal gyrus and IFS (104 electrodes, 18 patients). Right: Spectrogram and electrode placements for posterior superior temporal sulcus (pSTS) (electrodes: 46; patients: 21). (E): Exemplar electrode displaying greater broadband gamma activity (BGA) for limited narrowing trials in pSTS.

Trials with limited semantic narrowing revealed greater BGA in and around posterior superior temporal sulcus (pSTS) beginning approximately 250 ms after the onset of the final word, and lasting until approximately 900 ms (300–500 ms: *β* = 0.10 (SD: 0.04); *p* = 0.001; 500–700 ms: *β* = 0.08 (SD: 0.03); *p* < 0.001). Greater BGA for limited narrowing trials was also found in MPC (500–700 ms: *β* = 0.12 (SD: 0.07); *p* = 0.003), IFS (300–500 ms: *β* = 0.11 (SD: 0.05); *p* = 0.008; 500–700 ms: *β* = 0.12 (SD: 0.07); *p* = 0.004), and anterior temporal lobe (500–700 ms: *β* = 0.06 (SD: 0.01); *p* = 0.004).

## Discussion

Our evaluation of the spatiotemporal dynamics of orthographic sentence comprehension and the mapping of these representations onto common object referents identified distinct roles for closely adjacent loci in lateral inferior frontal and lateral posterior temporal cortex. Whilst frontal gyral structures are commonly implicated in the literature (Friederici, 2017; Matchin and Hickok, 2020), our intracranial recordings with depth electrodes also recorded from sulcal structures. The inferior frontal sulcus (IFS) was uniquely sensitive across these semantic processes, exhibiting a clear mosaic-like patchwork of activity across its sub-regions (Fig. 5). Although many models have proposed anterior-posterior distinctions for inferior frontal regions with respect to certain higher-order language functions (Hagoort, 2005; Hagoort and Indefrey, 2014; Kuhnke et al., 2017; Matchin and Hickok, 2020; Price, 2010), our intracranial recordings of distinct semantic effects provides a more complex picture. Though we indeed find support for separable effects in anterior and posterior inferior frontal gyrus (IFG) – semantic coherence for the former, semantic narrowing for the latter – there was also found to be some functional overlap; both sites were sensitive to the referential status of sentences. Neighboring sulcal loci also showed effects that were not only earlier than those in IFG, but also more topographically complex. Centered around IFS and spreading out dorsally towards middle frontal gyrus and ventrally towards anterior/posterior inferior frontal gyrus, we uncovered a cortical mosaic of functional specialization for reference, coherence and semantic narrowing.

**Figure 5:**
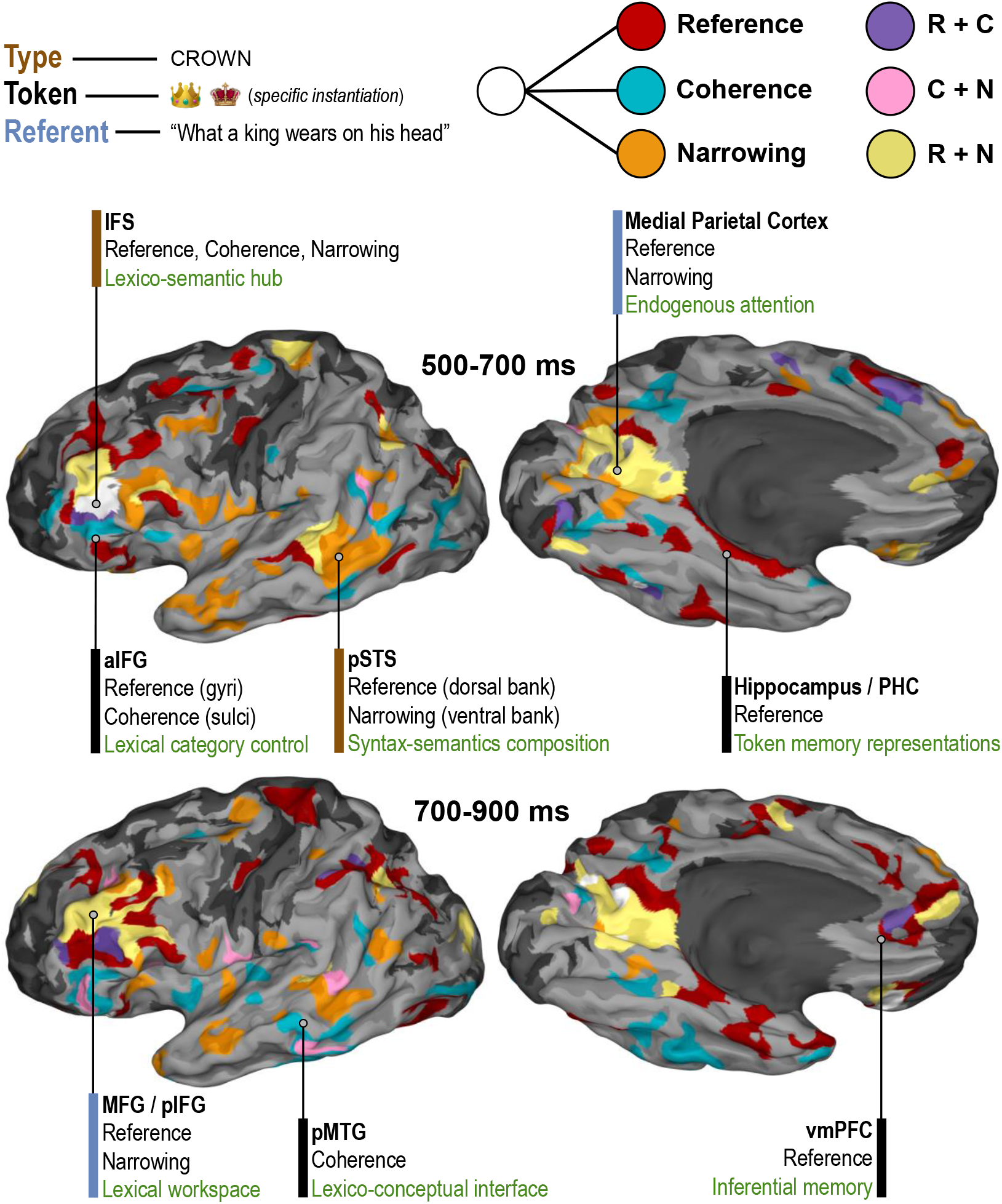
Summary model for distinct components of linguistic meaning. Model derived from present analyses of semantic reference (referential vs. non-referential sentence endings), semantic coherence (weak vs. strong violations of lexico-semantic rules for non-referential trials) and semantic narrowing (limited vs. strong narrowing). Top: Color scheme for Type, Token and Referent (matched to colored bars next to regional descriptions), and also for regions showing some conditional difference for Reference, Coherence and Narrowing (matched to colored clusters on brain plots), across two main time windows of interest. Bottom: Conjunction SB-MEMA depicting regions sensitive to conditional contrasts, regardless of directionality (threshold: %BGA > 5%, t > 1.96, patient coverage ≥ 2). Functional descriptions in green (bottom entries) are inferred from existing literature and are not intended to be monolithic, but specific to the present task of definitional object naming (see Discussion).

Early processing windows implicated IFS in all aspects of semantics, and in later windows sub-portions of IFS exposed a clearer functional tessellation with some remaining overlap.

Posterior superior temporal sulcus (pSTS) and posterior middle temporal gyrus (pMTG) also jointly contributed to all processes, with a portion of overlap in pSTS for reference and narrowing. The temporal progression of sensitivity in posterior temporal cortex had a clear delineation between early effects of narrowing, then reference, and finally coherence. Together, our recordings of posterior temporal and inferior frontal cortex provide a comprehensive, whole brain map of the semantics system. Portions of fronto-temporal cortex were engaged for all aspects of language semantics, but become rapidly dedicated to certain processes at distinct times. Previous intracranial work has implicated pSTS in the initial construction of meaningful phrases, and IFS in the later evaluation of these phrases in the service of phrase-picture matching (Murphy et al., 2022b). Our results further highlight the role of pSTS-IFS in semantic composition demands, with early narrowing effects in pSTS (onset ∼ 250 ms) being followed by effects for all compositional processes in IFS (∼ 500 ms) (Fig. 5).

We also found that medial parietal cortex (MPC), hippocampus, parahippocampal gyrus (PHG), and ventro-medial prefrontal cortex (vmPFC) are engaged during the processing of sentences that referred to common objects. We found greater activity in posterior superior temporal gyrus and anterior inferior frontal gyrus (aIFG) for semantic violations in non-referential contexts (Fig. 3). These findings have direct relevance to models of semantics and dysnomia, and we discuss here the most salient implications.

The network implicated in the ability to successfully generate names of visually presented common objects consists of the inferior frontal cortex, mid-fusiform cortex and the supplementary motor area (SMA) (Conner et al., 2014; Forseth et al., 2018; Swanson et al., 2020). Our results provide additional details about particular integrative components of semantic processing, with our analyses dissociating semantic narrowing and coherence. We also reported on the basic build-up of local cortical activity across normal sentence processing (Fig. 2). Examining basic contrasts for orthographic sentence processing (Fig. 2), we can see regions more likely involved in lexical access from the Final Word – Word 1 contrast, and regions more likely involved in semantic integration from the Final Word – Penultimate Word contrast. The major nodes we identified in this network are likely responsible for distinct components of semantic integration (Poeppel et al., 2012), with areas such as pMTG, posterior cingulate and vmPFC becoming significantly active only later in the sentence. As such, processes relating to situation model construction may be subserved by these regions, with vmPFC being implicated recently in associative inference and memory integration (Spalding et al., 2018).

### Medial temporo-parietal involvement in reference

Previous research into linguistic reference to both visual objects and sounds implicated greater engagement of MPC from 500–600 ms after the orthographic presentation of the referential word in a sentence (Brodbeck et al., 2016). These regions form part of the default network, which is engaged during endogenous attentional tasks, with representational search likely being greater for sentences permitting successful selection of objects from the lexicon (Buckner and DiNicola, 2019; Smallwood et al., 2021). Default network activity is related to processing rich representations of events, either real or imagined (Hassabis and Maguire, 2009), of the kind inferred by patients in our task. Relatedly, our effects in dorsal frontal sites seem to overlap with a node in the multiple demand network (Assem et al., 2020; Duncan et al., 2020; Fedorenko et al., 2013), with has been implicated in linguistic processing difficulty, seemingly congruent with our semantic narrowing effects in IFS and MFG.

Linguistic reference to coherent entities has been shown to implicate medial frontal and medial parietal regions, along with classic perisylvian language regions (Ferstl et al., 2008). Sentence pairs introducing a conjoined subject (e.g., “John and Mary”) result in greater MPC engagement compared to individual referents (Boiteau et al., 2014). Sentences containing referentially ambiguous pronouns (e.g., “Saul told Mike that he… “) have been shown to result in medial parietal activity (Nieuwland et al., 2007). Our findings concerning medial parietal regions seem in accord with the involvement of the MPC in the posterior medial system (Ranganath and Ritchey, 2012), which is part of the distributed memory network and is involved in context memory, constructing situation models, and combining concepts for distinct categorical domains (Burianova and Grady, 2007; Burianova et al., 2010; Lundstrom et al., 2005; Maguire, 2001; Rabini et al., 2021; Rajah and McIntosh, 2005; Schacter and Addis, 2007; Zwaan, 2016). The MPC has been shown to contain selective regions for the recall of information about familiar people and places, but does not demonstrate activity during visual object naming (Peer et al., 2015; Silson et al., 2019; Woolnough et al., 2020), and the precuneus has been implicated in the encoding of complex memories and actions (Hebscher et al., 2019; Rolls, 2019).

In addition to MPC, the posterior medial system incorporates the hippocampus and PHG, another region observed in our study. Hippocampal theta tracks the amount of contextual linguistic information in a sentence, such that in predictable sentence contexts theta power increases during sentence presentation (Piai et al., 2016). Other recent intracranial work has indicated a role for medial temporal lobe in verbal working memory (Boran et al., 2020). This suggests that portions of hippocampus and PHG actively relate incoming words to sentence contexts and contribute memory-related representations needed to successfully search the (narrowed) lexicon. We discovered early hippocampal theta power increases for referential trials (100–500 ms) followed directly by BGA increases in PHG (300–1000 ms); it seems plausible that joint MPC-hippocampal engagement underlies contextual associations (Aminoff et al., 2013; Benítez-Burraco and Murphy, 2019; Murphy et al., 2022a).

Our results are also concordant with the findings that hippocampal-complex damage leads to impairments in semantic association tasks testing words learned prior to damage (Klooster and Duff, 2015), and that hippocampal theta power increases during episodic memory retrieval (Lega et al., 2012). It has been argued that the hippocampus might contribute to language’s facility for displacement (Benítez-Burraco, 2021; Corballis, 2019), or the capacity to refer to objects/events outside current spatiotemporal context, and also the tracking of situation/context boundaries (Maurer and Nadel, 2021), potentially concordant with its involvement in definitional object reference.

### Semantic coherence in the absence of lexical access from definitions

Some of our stimuli were semantically coherent, but did not refer to a common object. These non-referential sentences (weak violations) resulted in greater BGA in aIFG, IFS, angular gyrus and pMTG. The early effect for weak semantic violations in aIFG (onset around 300 ms) possibly indexes the entry into the syntactic workspace of the final word and the successful wrap-up effect for semantically legal structures, and the subsequent control of the appropriate lexical category (Matchin and Hickok, 2020). Previous work has implicated IFG in various types of semantic information during different tasks such as plausibility and acceptability judgements (Friederici, 2011; Hagoort and Indefrey, 2014). Our results are generally in line with the role of IFG in effortful lexico-semantic processing (Branco et al., 2020), but afford a more nuanced picture: Lateral aIFG exhibits greater BGA for non-referential relative to referential sentences, and neighboring aIFG and IFS show greater BGA for semantically coherent non-referential sentences, relative to incoherent non-referential sentences. This leads to a fine-grained spatiotemporal map of anterior IFG activity. Lateral aIFG activity marks semantic violations and cortex abutting IFS responds to semantic coherence. We note that during basic sentence comprehension (Fig. 2) these portions in aIFG and IFS are engaged jointly, suggestive of coordinated semantic processes, dissociable with fine spatiotemporal resolution and appropriate behavioral contrasts isolating coherence, reference and semantic narrowing.

Lastly, we note that inferior parietal regions (angular gyrus) were only implicated in this specific contrast for semantic coherence, but no other contrasts. This region was also only moderately active towards the end of sentence comprehension (Fig. 1). Though not as engaged as other regions, these results support suggestions that this region codes for event/thematic structure (Binder and Desai, 2011; Matchin and Hickok, 2020).

### Narrowing the lexical search space

Trials that exhibited a limited degree of semantic narrowing (i.e., that did not clearly reduce the space of possible lexical items to select for articulation) resulted in greater recruitment of IFS, pSTS, MPC, and ATL. Recruitment of pSTS was earliest (approximately 200 ms), and may pertain to composition-related demands or lexical search effort (Fig. 5–7), indexing greater engagement of compositional processing due to semantic narrowing demands (Flick and Pylkkänen, 2020; Matchin and Hickok, 2020), in particular given previous intracranial work implicating this region in basic semantic composition and lexicality (Murphy et al., 2022b). ATL engagement is in line with its apparent role in conceptual processing and entity-related (e.g., common object) representations (Thye et al., 2021; Turken and Dronkers, 2011).

While medial brain structures often get overlooked in major models of higher-order language, our results suggest that a more extended network is involved in elementary components of sentence comprehension and lexical access. We provide a general model for the neural representation of linguistic meaning derived from a conjunction map of the above analyses (Fig. 5), highlighting also general engagement in processes pertaining to higher-order semantics.

Our results can also be viewed in the context of the three core components of linguistic meaning – types, tokens, referents – that seem to more significantly implicate, respectively, left temporal, inferior frontal, and inferior parietal cortices (Baggio, 2018). A type is a general category of an entity (e.g., FLOWER), a token is a particular concrete instantiation (e.g., a specific flower), and a referent arises via the explicit denotation of a token (e.g., “That yellow thing in the garden”). Our paradigm involved descriptions of referents calling upon conceptions of general types. Although our paradigm was not designed to specifically adjudicate between these three aspects, the immediate compatibility here appears to be between our main parietal effects and the connections with processing the discourse referent. MPC is sensitive to referential and semantic coherence, and may form part of the lexical search process and the construction of *referents*; we note that although this region is involved in general category sensitivity (e.g., faces, scenes), strictly within the context of our task its involvement seems centered on establishing discourse referents. A small portion was sensitive to all conditions during later windows, but the dominent response profile was specialized. Given the role of (para)hippocampus in memory, the activity differences we found exclusively for the referential/non-referential contrast may index referent-specific memory traces and hence *tokens* (Fig. 7). The sensitivity of pSTS to semantic narrowing and referential violations (separately and jointly, depending on sub-region), and the sensitivity of IFS to all semantic processes, point towards an involvement in semantic *types*. More generally, it may be possible to adjudicate here between strictly lexico-semantic engagement (pSTS, IFS) and more domain-general semantic memory activation (dorsal frontal, medial parietal, ventral temporal) (Binder and Fernandino, 2015; Hagoort, 2020), in particular given that parahippocampal cortex, precuneus and vmPFC are all implicated in general semantic processing (Binder et al., 2009). Overall, this work suggests that current models apportioning clear frontal, temporal and parietal separation between semantic representations (e.g., types, tokens and referents) (Baggio, 2018) need to accommodate the richly interwoven, mosaic-like architecture of the natural language semantics system.

### Summary

We reported extensive whole brain intracranial mapping of semantically coherent orthographic sentence representations, providing insights into conceptual access. We reported diverse roles for posterior temporal and inferior frontal language regions, and dissociated regional contributions to distinct semantic processes. In particular, our intracranial recordings afforded access to sulcal structures as well as the lateral surface of the cortical mantle, with IFS uniquely responding to all semantic processes, potentially indexing its role as a higher-order lexico-semantic hub. Rather than finding posterior temporal and inferior frontal engagement only for successful semantic integration, or only for semantic violation detection, we found that partially overlapping sub-regions of these loci are engaged in both (within pSTS, IFS and aIFG). This topography of functional arrangement implies the existence of small scale regional networks in the lateral frontal and temporal cortex that then connect with the larger scale semantic network distributed across lobes. A clearer understanding of these systems will pave the way for deeper insight into developmental dyslexia, acquired language impairment and potentially enable better neuro-rehabilitative and neuroprosthetic approaches for these disorders.

## Materials and Methods

### Participants

58 patients (11 male, 18-41 ± 5.7 years, IQ 95 ± 15, 2 left-handed) participated in the experiment after written informed consent was obtained. All were native English speakers. Participants with significant additional neurological history (e.g., previous resections, MR imaging abnormalities such as malformations or hypoplasia, or those with prosopagnosia) were excluded. All experimental procedures were reviewed and approved by the Committee for the Protection of Human Subjects (CPHS) of the University of Texas Health Science Center at Houston as Protocol Number: HSC-MS-06-0385.

### Electrode Implantation and Data Recording

Data were acquired from either stereotactically placed depth electrodes (sEEGs; 56 patients) or subdural grid electrodes (SDEs; 2 patients). SDEs were subdural platinum-iridium electrodes embedded in a silicone elastomer sheet (PMT Corporation; top-hat design; 3mm diameter cortical contact), and were surgically implanted via a craniotomy (Conner et al., 2011; Pieters et al., 2013; Tandon, 2012; Tong et al., 2020). sEEGs were implanted using a Robotic Surgical Assistant (ROSA; Medtech, Montpellier, France) (Rollo et al., 2020; Tandon et al., 2019). Each sEEG probe (PMT corporation, Chanhassen, Minnesota) was 0.8 mm in diameter and had 8-16 electrode contacts. Each contact was a platinum-iridium cylinder, 2.0 mm in length and separated from the adjacent contact by 1.5-2.43 mm. Each patient had 12-20 such probes implanted (total electrode count: 11,328; M = 199). Following surgical implantation, electrodes were localized by co-registration of pre-operative anatomical 3T MRI and post-operative CT scans in AFNI (Cox, 1996). Electrode positions were projected onto a cortical surface model generated in FreeSurfer (Dale et al., 1999), and displayed on the cortical surface model for visualization (Pieters et al., 2013). Intracranial data were collected during research experiments starting on the first day after electrode implantation for sEEGs and two days after implantation for SDEs. Data were digitized at 2 kHz using the NeuroPort recording system (Blackrock Microsystems, Salt Lake City, Utah), imported into Matlab, initially referenced to the white matter channel used as a reference for the clinical acquisition system and visually inspected for line noise, artifacts and epileptic activity. Electrodes with excessive line noise or localized to sites of seizure onset were excluded. Each electrode was re-referenced to the common average of the remaining channels. Trials contaminated by inter-ictal epileptic spikes and trials in which participants responded incorrectly were discarded.

### Stimuli and Experimental Design

Patients were asked to quickly and accurately articulate the name of common objects in response to orthographic descriptions (mean: 6.5 words, range: 3-12 words). Sentences varied in length (3–12 words) to reduce the predictability of the location of the final, definition-determining word. A fixation cross was presented in the centre of the screen for 700ms, followed by each successive word in the sentence, each appearing for 500ms, and then a blank screen was presented for 1 second. Patients had 2 seconds to respond. The number of trials per block across the full experiment was as follows: Referential (42), Non-Referential (42). Most patients undertook two blocks (n=48), with some only completing a single block (n=10). Stimuli were presented using Psychtoolbox (Kleiner et al., 2007) on a 15.4” LCD screen positioned at eye-level, 2-3’ from the patient. Auditory stimuli were presented using stereo speakers (44.1 kHz, MacBook Pro 2008).

To determine the profile of semantic narrowing effects in our stimuli, we conducted a norming study on a non-clinical population (n = 80, 50 female, mean age: 34, range: 18–71). All were native English speakers, and had a 90% minimum approval rating on Prolific Academic from which they were recruited to take part in the questionnaire (run via the Qualtrics online platform). Participants were paid 10 USD per hour, and the average completion time was 26 minutes. Participants were presented with the referential trials from the main experiment, word-by-word, and were asked to type any possible corresponding words that might match the ongoing description upon the presentation of a new word. For example, they were first shown “A”, followed by “A round”, then “A round red”, and finally “A round red fruit”. Most participants answered “apple” after being presented with “A round red”, with many answering “ball” or “circle” after seeing “A round”.

### Signal Analysis

A total of 13,298 electrode contacts were implanted in these patients; 9,388 of these were included for analysis after excluding channels proximal to the seizure onset zone or exhibiting excessive inter-ictal spikes or line noise. Analyses were performed by first bandpass filtering the raw data of each electrode into broadband gamma activity (BGA; 70-150 Hz) following removal of line noise and its harmonics (zero-phase 2nd order Butterworth band-stop filters). A frequency domain bandpass Hilbert transform (paired sigmoid flanks with half-width 1.5 Hz) was applied and the analytic amplitude was smoothed (Savitzky-Golay FIR, 3rd order, frame length of 251ms; Matlab 2019b, Mathworks, Natick, MA). BGA was defined as percentage change from baseline level; -500ms to -100ms before the presentation of the first word in each trial. Periods of significant activation were tested using a one-tailed t-test at each time point and were corrected for multiple comparisons with a Benjamini-Hochberg false detection rate (FDR) corrected threshold of q<0.05. For the grouped analysis, all electrodes were averaged within each subject and then the between-subject averages were used.

To provide statistically robust and topologically precise estimates of BGA, and to account for variations in sampling density, population-level representations were created using surface-based mixed-effects multilevel analysis (SB-MEMA) (Conner et al., 2011; Fischl et al., 1999; Kadipasaoglu et al., 2014, 2015). This method accounts for sparse sampling, outlier inferences, as well as intra- and inter-subject variability to produce population maps of cortical activity. A geodesic Gaussian smoothing filter (3 mm full-width at half-maximum) was applied. The minimum criterion for the family-wise error rate was determined by white-noise clustering analysis (Monte Carlo simulations, 5000 iterations) of data with the same dimension and smoothness as that analyzed (Kadipasaoglu et al., 2014). Results were further restricted to regions with at least three patients contributing to coverage. Due to the proximity to loci with typical eye movement artefacts (Kovach et al., 2011), we also evaluated right-hemispheric ventromedial prefrontal cortex and verified that reported effects were specific to the left hemisphere (Suppl. Fig. 1). Likewise, we confirmed that only the language-dominant left hemisphere medial structures, and not right hemisphere structures, index sensitivity to referential sentences.

Our regions of interest were clustered from a number of Human Connectome Project (HCP) regions (Glasser et al., 2016): Medial parietal cortex (RSC, POS1, POS2, v23ab, 7m, 31pv, 31pd, d23ab); hippocampus and parahippocampal gyrus (PreS, EC, PHA1); ventromedial prefrontal cortex (a24, p32, 10r, s32, 10v, 25, OFC); middle frontal gyrus (8C, 46, p9, 46v); dorsal inferior frontal gyrus and inferior frontal sulcus (IFSa); posterior superior temporal sulcus (TPOJ1, TPOJ2). For the low frequency analysis in hippocampus, we manually checked electrode placements to ensure a distinction between electrodes placed within hippocampus proper and within neighboring parahippocampal sites (Parahippocampus = PreSubiculum/Entorhinal cortex/Parahippocampal Area 1).

The language system is richly interwoven with episodic memory networks, with segregation between these systems resulting more from investigator domain expertise than neurobiological underpinnigs. Given that hippocampal theta is modulated by sentence predictability (Piai et al., 2016), theta phase-coupling occurs between hippocampus and left superior temporal gyrus increases during correct (vs. incorrect) sentences (Pu et al., 2020), and theta power in inferior frontal cortex and central EEG sites is linked to language comprehension (Hald et al., 2006; Mellem et al., 2013; Momsen and Abel, 2021), we also examined low frequency dynamics across these sites.

Anatomical groups of electrodes were delineated, firstly, through indexing electrodes to the closest node on the standardized cortical surface (Saad and Reynolds, 2012), and secondly, through grouping channels into parcellations determined by Human Connectome Project (HCP) space (Glasser et al., 2016). Parametric statistics were used since HCP regions of interest contained >30 electrodes. To determine significant activity increases from baseline, a two-sided paired t-test was evaluated at each time point for each region and significance levels were computed at a corrected alpha-level of 0.05 using an FDR correction for multiple comparisons. To determine a significant difference between conditions in the activation of a region, the cumulative BGA in a specified time window was evaluated with a two-sided paired t-test at an alpha-level of 0.01.

## Acknowledgements

We express our gratitude to all the patients who participated in this study; the neurologists at the Texas Comprehensive Epilepsy Program who participated in the care of these patients; and the nurses and technicians in the Epilepsy Monitoring Unit at Memorial Hermann Hospital who helped make this research possible. We also thank Kathryn Snyder and Oscar Woolnough for feedback on earlier drafts. This work was supported by the National Institute of Neurological Disorders and Stroke NS098981. Human subjects: Patients participated in the experiments after written informed consent was obtained. All experimental procedures were reviewed and approved by the Committee for the Protection of Human Subjects (CPHS) of the University of Texas Health Science Center at Houston as Protocol Number: HSC-MS-06-0385.

## Author Contributions

NT, KJF, CD designed research; EM, KJF, CD, PSR performed research; EM analyzed data; EM wrote the first draft of the paper; EM, NT edited the paper; NT conducted funding acquisition and neurosurgical procedures.

## Data and Code Availability

The datasets generated from this research are not publicly available due to them containing information non-compliant with HIPAA and the human participants the data were collected from have not consented to their public release. However, they are available on request from the corresponding author. The custom code that supports the findings of this study is available from the corresponding author on request.

## Supplementary Material

**Supplementary Figure 1:**
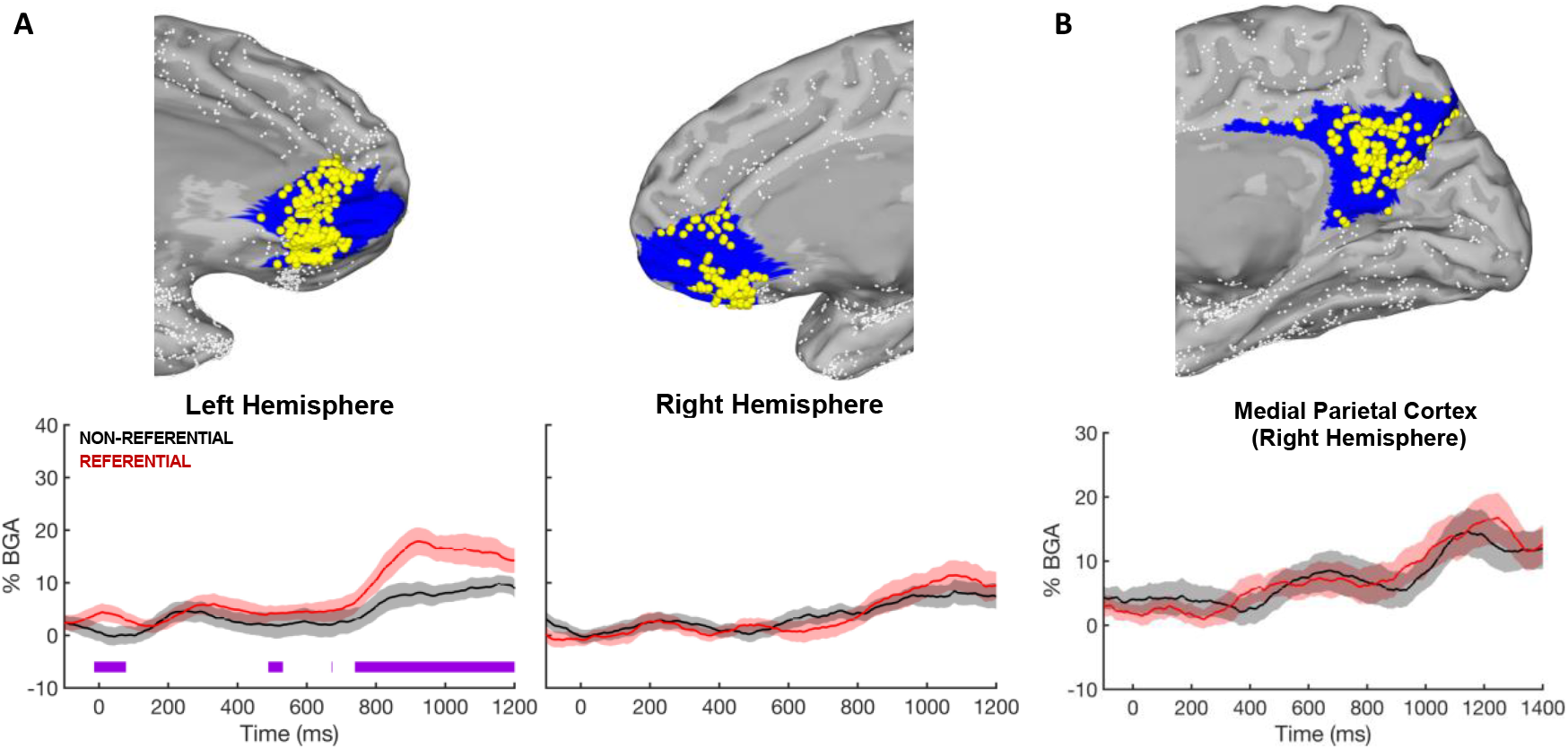
Hemispheric contrasts of reference. (A): Electrodes placed within left vmPFC (233) and their corresponding BGA trace, alongside the right hemisphere (126). FDR-corrected significance bars plotted in purple. Time-locked to final word onset. (B): Electrodes placed within the right medial parietal cortex (139) from patients with bilateral coverage (31) and their corresponding BGA traces (for left hemisphere, see Figure 3).

## Notes

**Conflicts of interest:** The authors declare no competing interests.

### Competing Interest Statement

The authors have declared no competing interest.

