## Supplementary figures and images for "The spatiotemporal dynamics of semantic integration in the human brain"

### Supplementary Figure 1

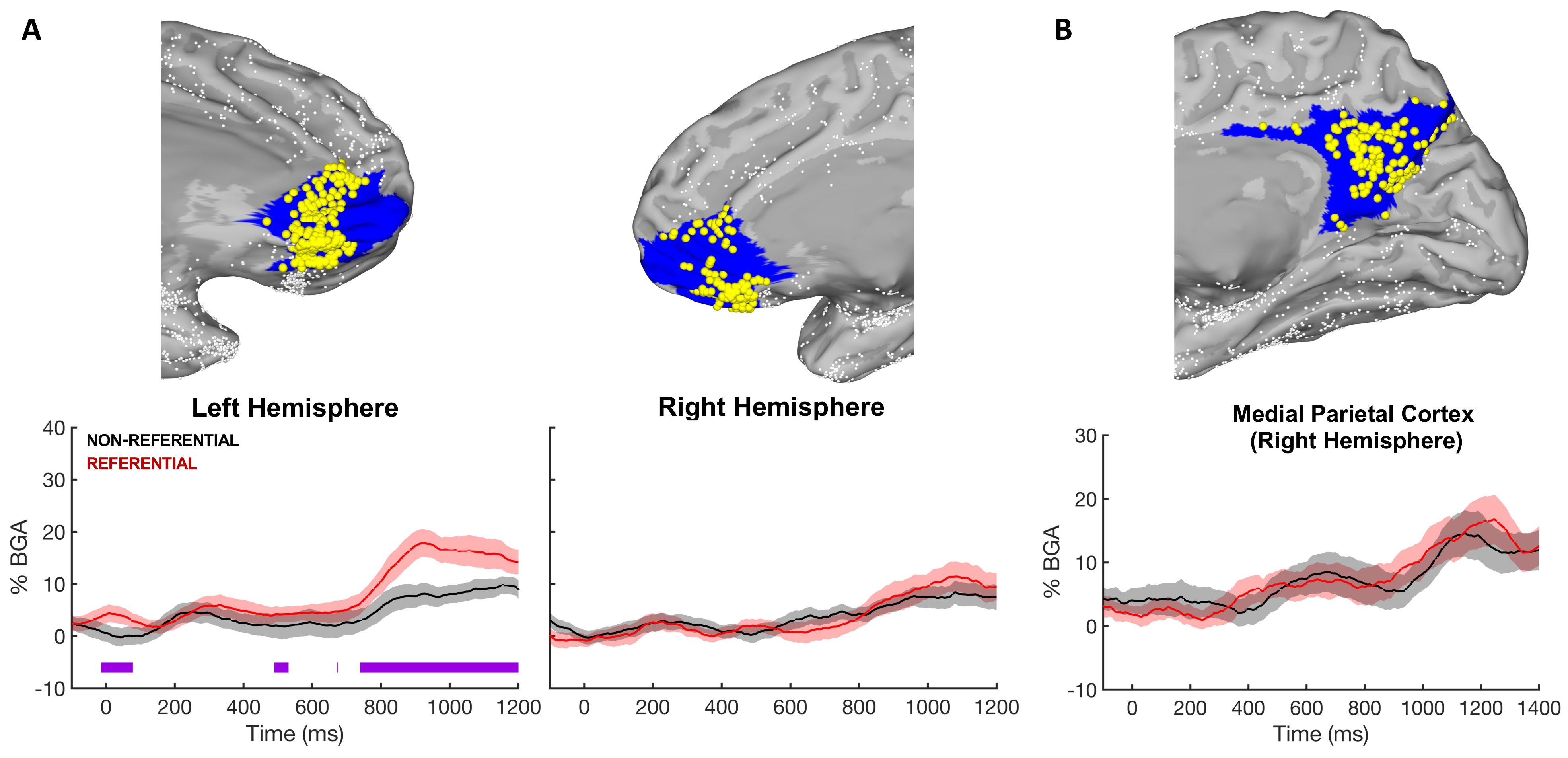
